# The genome assembly and detection of biosynthetic gene clusters among four novel microbial isolates from deep subsurface rock biofilms

**DOI:** 10.1101/2025.09.13.675994

**Authors:** Diing D.M Agany, Jessica LS Zylla, Shiva Aryal, Oxana Gorbatenko, David Bergmann, Venkataramana Gadhamshetty, Etienne Z. Gnimpieba

**Affiliations:** Biomedical Engineering, University of South Dakota, Sioux Falls, SD 57107, USA; Black Hills State University, Spearfish, SD, 57799, USA; Department of Civil and Environmental Engineering, South Dakota School of Mines and Technology, Rapid City, SD 57701, USA; 2Dimensional Materials for Biofilm Engineering Science and Technology (2DBEST) Center, South Dakota School of Mines and Technology, Rapid City, SD 57701, USA

**Keywords:** Whole Genome Assembly, Old mine tunnel, Subsurface rock, Biofilms, Deep biospheres, Cave silver, Biosynthetic Gene Clusters, secondary metabolites, Natural products

## Abstract

Tunnels in deep underground mines provide a unique interface between the surface and deep subsurface habitats. There, microbes from the surface may enter the deep tunnels as surface air is drawn into the deep mine tunnels to provide ventilation. This extreme hosting environment provides a condition for microbes to develop novel capabilities, such as the production of natural products of biotechnology or medicinal importance. This study characterized the genomes of four novel isolates from deep subsurface biofilms of the previously abandoned gold mine, which was used as a model for the underground study. Here the microbiome samples were obtained from thin, whitish, glistening biofilm samples naturally formed on the rock walls (1478 meters deep at SURF). These samples are herein referred to as “cave silver” biofilms. The samples provide 10 GB of high-quality whole genome sequences that were assembled into contigs/scaffolds and structurally and functionally annotated against various databases. Subsequently, the genomes were analyzed for biosynthetic gene clusters (BGCs) encoding secondary metabolites of biotechnology and medical importance using the antiSMASH and the NaPDoS web servers, respectively. In brief, the assemblies produced four drafted genomes of different lengths and annotated features for each strain’s genome, including gene clusters involved in the quorum sensing (QS) pathway. Several BGC-encoded secondary metabolites of natural products or compounds such as polyketides (PKS, PKS III, ketosynthase domain-KS), non-ribosomal peptides (condensation domain, NRPS), and terpenoids (terpenes I, polyenes type II), were identified. Furthermore, many overexpressed enzymes, most of which are shared among the strains and some unique to individual bacteria strains, were identified, revealing possible shared and individualized strain activities in the biofilms during the sample collection. CRISPR peptides were also detected in three of the four bacteria strains. In this study, we examine the genomes of microbes colonizing extreme subsurface environments and how these conditions promote the development of microbial systems that are potentially capable of producing medicinally natural products and secondary metabolites that may be used to enhance advancements in biotechnology products.

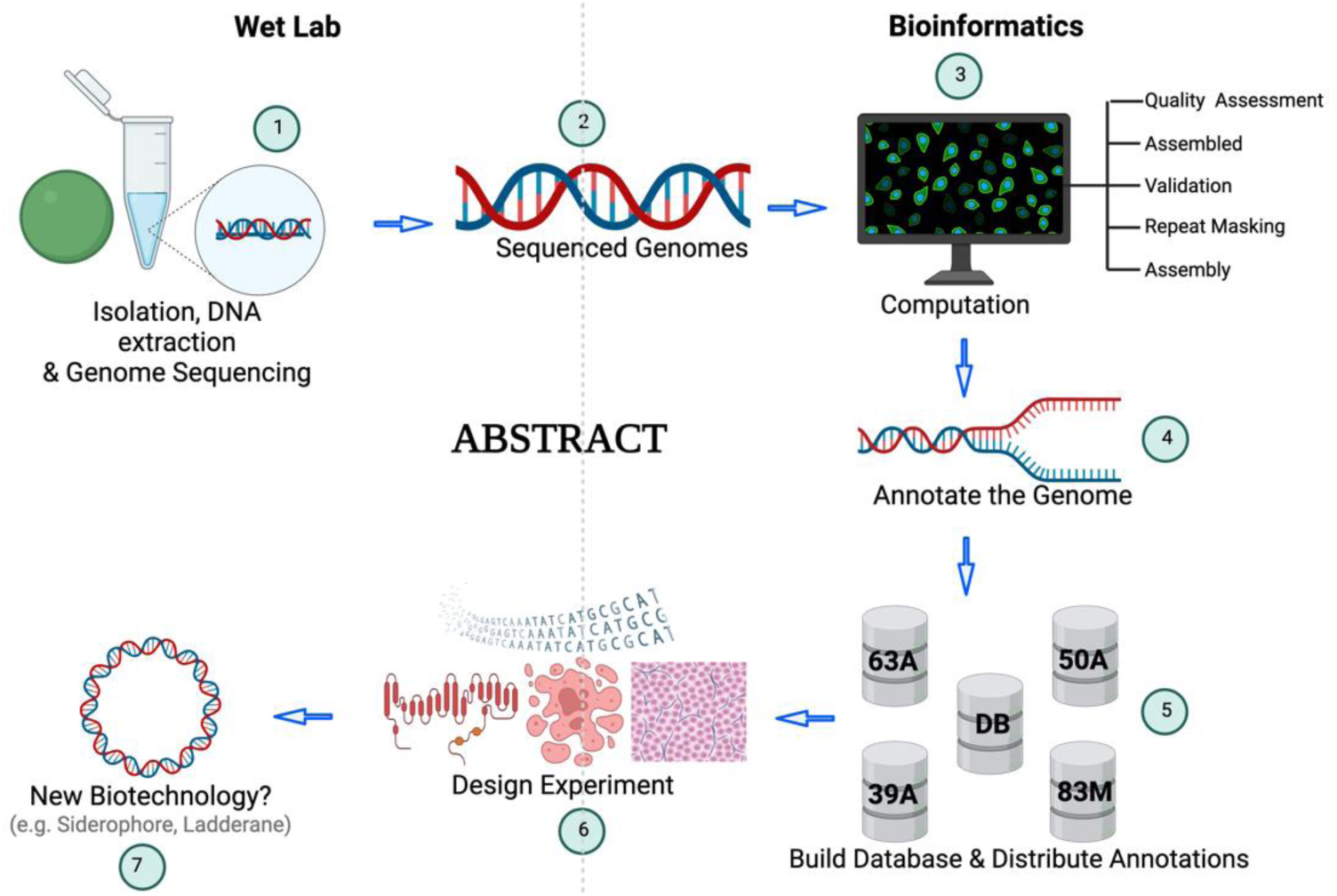

**Importance:** Deep subsurface environments host unique microbial communities with specialized adaptations. By characterizing genomes from “cave silver” biofilms, this study reveals biosynthetic gene clusters and pathways for secondary metabolite production, including polyketides, peptides, and terpenoids. These findings highlight the potential of extreme-environment microbes as a novel source of biotechnologically and medically valuable natural products.

## Introduction

The naturally hostile conditions of the extreme environments (hot, dark, and deep subsurface) constantly exert evolutionary pressures on the microbiome. According to Singh et al, such environments are found in deep subsurface like old mines and are favorable for finding hardy novel microbes (Singh, N. K. 2017). The deep underground biosphere consists of rocks and aquifers extending over 2 km beneath the earth’s surface. In recent years, there has been an increased interest in sampling and analysis of deep subsurface sites (Borgonie, G., 2015) to characterize the composition and diversity of microbial communities (Dong, H. et al. 2018, Drake, H. et al. 2017) as well as their function and interaction. For example, Nyyssönen et al. (Nyyssönen, M. et al. 2012) studied sulfate-reducing bacteria and methanogenic Archaea 2516 meters deep in the Finnish crystalline bedrock aquifers.

With the increased depth of deep biosphere sampling capabilities and the availability of the next generation sequencing (NGS) technology, microbes that could not be reached for sampling are now isolated and sequenced or directly sequenced to study their genomes and the niche ecosystems (Rinke, C. et al. 2013). The Sanford Underground Research Facility (SURF); a former Homestake gold mine in Lead, South Dakota, USA (https://fanniesanfordlab.org/feature/sanford-underground-research-facility-overview) is an example of that environment. The tunnels at deep underground mines, such as the SURF, create a special connection between the surface and deep underground areas. Here, warm, oxygen-free rocks and water meet the cooler, oxygen-rich air in the mine’s tunnels, and microbes from the surface may enter the deep tunnels as surface air is drawn into the deep mine tunnels to provide ventilation.

In one tunnel on the 1478 deep level of SURF, warm, moist air from the distal portions of the tunnel meets with cooler air, which is drawn down shafts during ventilation, resulting in a region of condensation on the tunnel walls. As a result, the rock walls support extensive thin, whitish, glistening biofilms resembling the “cave silver” biofilms found in some limestone caves (Pašic, L. et al. 2010) and lava tube caves (Lavoie, K.H. et al. 2017) in outward appearance and structure.

The cave silver biofilms in SURF, similar to these found in cave silver elsewhere, contain abundant mycelial bacteria from the family Pseudomonadaceae. However, 16S rRNA gene sequences indicate that nearly all operational taxonomic units (at the 97% level of similarity) present in SURF cave silver are absent in limestone or lava tube caves (Thompson, E. et al. 2020).

To study the microbial inhabitants of this extreme environment, Brar and Bergmann (Brar et al. 2019) applied culture-based techniques to examine the cave silver biofilm bacteria communities in the SURF. Their study obtained various isolates, primarily consisting of Alpha-Proteobacteria and Actinobacteria superfamilies, some of which were genera not previously cultured. These novel isolates were chosen for whole-genome sequencing to confirm their identities and study their genomes. Among them, collected at 1478 meters depth (SURF) and confirmed by 16S and this sequencing study, are four isolated genera of *Allorhizocola sp*, *Taonella* sp., *Dongia* sp., and *Pseudonocardia* sp.

Using the sequence data, each of the genus genomes was assembled, annotated, and analyzed for biosynthetic gene cluster (BGC) presence and their encoded secondary metabolites. The genomes were examined to further understand the life of microbes under extreme conditions, including their utility of carbon sources, byproducts, and adaptation to an obligate symbiotic lifestyle (Beam, J. P. et al. 2020).

## Materials and Methods

### Bacterium collection, isolation, and culturing

Bacteria were collected from “cave silver” biofilms on phyllite rocks lining a tunnel on the 17 Ledge of SURF and were cultured as described by Brar and Bergmann (Brar A.K. and Bergmann D., 2019). Four isolates were chosen for genome sequencing: (1) an isolate (strain 63A) identified as *Pseudonocardia*, (2) an isolate belonging to a novel genus of Alpha-Proteobacteria (strain 50A) related to *Dongia*, (3) another isolate (strain 39A) of a novel genus belonging to the Alpha-Proteobacteriaceae related to *Taonella*, and (4) an isolate (strain 83M) identified as *Allorhizocola sp.* (**Table 1**) based on 16S rRNA gene sequences. Throughout the paper, the authors refer to these isolates as 63A, 50A, 39A, and 83M.

**Table 1:** Study Name: SURF-bacteria isolates-33890863.

| IMG GENOME ID | 2830766779 | 2831136563 | 2831124213 | 2831130049 |
| --- | --- | --- | --- | --- |
| GENOME NAME | 63A | 50A | 39A | 83M |
| GENOME SIZE IN BP | 10321584 | 5115127 | 5738972 | 6404555 |
| GENE COUNTS | 10335 | 4987 | 5835 | 6513 |
| CDS COUNTS | 10234 | 4905 | 5751 | 6403 |
| RNA COUNTS | 86 | 68 | 71 | 99 |
| GC % | 68.6 | 66.45 | 68.13 | 66.94 |
| # OF SCAFFOLDS | 581 | 52 | 48 | 58 |
| # OF BPS | 10314156 | 5114878 | 5738717 | 6403083 |
| LEN SHORTEST SEQ | 150 | 249 | 255 | 206 |
| LEN LONGEST SEQ | 373151 | 458845 | 682729 | 769974 |
| AVG LENGTH | 17752 | 98363 | 119557 | 110398 |
| MEDIAN LENGTH | 1821 | 58886 | 42197 | 45127 |

### Genomic DNA extraction, library preparation, and sequencing

Genomic DNA from the isolates (63A, 50A, 39A, 83M) was extracted using the DNeasy Blood and Tissue Kit (Qiagen Inc., Waltham, MA 02452, USA), following the protocol for Gram-positive bacteria. All DNA was additionally purified using the Zymo Research DNA Clean and Concentrator-5 (Zymo Research, Irvine, CA, USA). Finally, DNA quantity was assessed using Qubit fluorometer 2 (Invitrogen™, Carlsbad, CA). In brief, the four libraries were constructed from extracted DNA using the Nextera XT DNA Library Preparation Kit (Illumina, San Diego, CA), following the manufacturer’s reference guide. Library quantities were assessed using a Qubit Fluorometer 2 (Invitrogen™, Carlsbad, CA) and an HT DNA HS reagent kit on a Lab Chip GX bioanalyzer (Caliper Life Sciences, Hopkinton, MA). Sequencing was done by the WestCore sequencing facility (Spearfish, SD) using a MiSeq instrument (Illumina, San Diego, CA). The libraries were pooled in equimolar amounts, and over 48 M (PE) paired-end reads 300 bp in length were generated using MiSeq Reagent Kit v3 (600-cycle) (Illumina, San Diego, CA). The raw read’s sequence quality was determined using FastQC v.11 (Andrews, S. 2010), configured inside the Blast2Go suite (Götz S. et al. 2008), and reads trimmed using FastP v.1.2 (Chen, S. et al. 2018), as shown in **Figures 1A and 1B**, which show the quality statistic for both forward and reversed short reads (83M). The other three strains showed a similar quality pattern and are not further reported here to reduce redundancy and inconvenience. Furthermore, pre *de novo* assembly, an additional primary statistical quality distribution of the raw reads was assessed using the CLC-Bio genome workbench. In these processes, quality assessments include the GC% contents, ambiguous base content, PHRED quality score distribution, nucleotide contributions, and sequence duplication levels. Additionally, the average quality statistics for each raw read were calculated, and low-quality bases were trimmed as necessary.

**Figure 1A:**
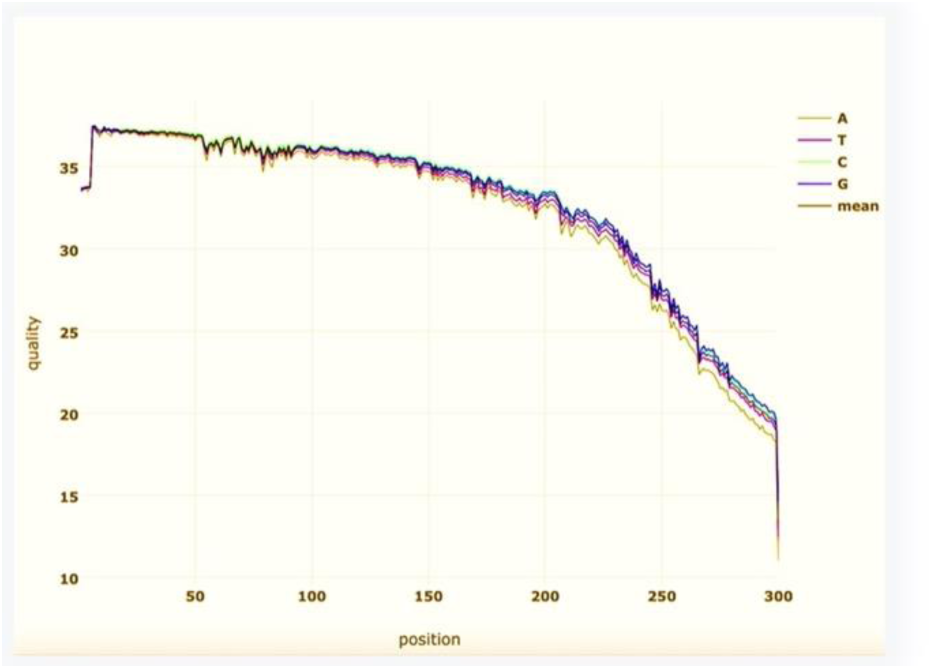
Untrimmed Sequences Quality score: a Phred score of sequences in a fast file showing the low-quality score of 12 before trimming.

**Figure 1B:**
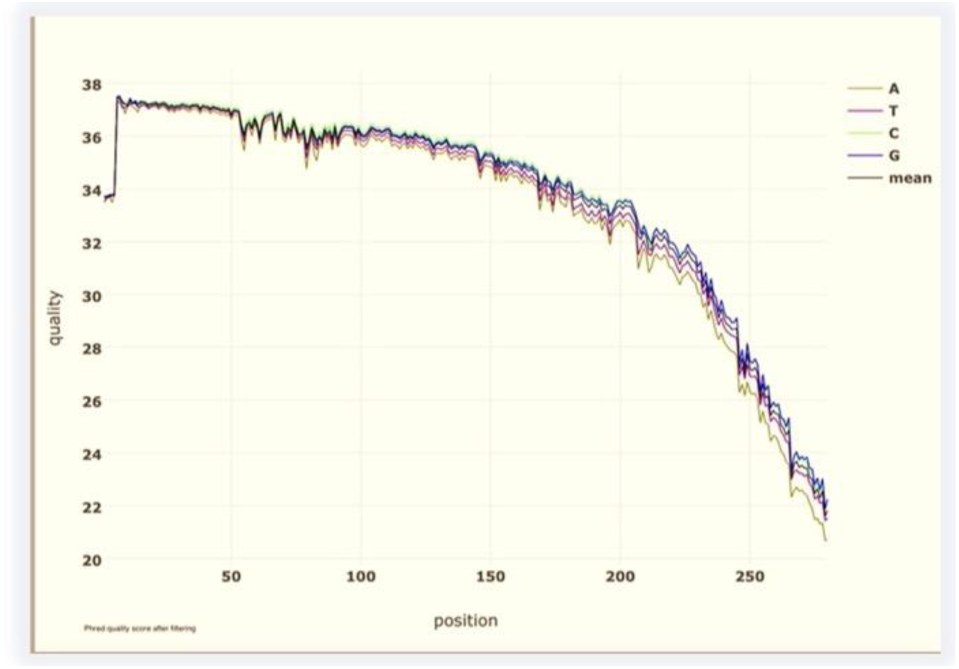
Trimmed Sequences Quality score: a Phred score of sequences in fastq file showing an improved quality score of 20 after trimming and removing low-quality sequences.

### Sequence assembly and genome analysis

Cleaned high-quality raw reads for each organism were *de novo* assembled using CLC-bio genome workbench (version 20.40). The quality of the de novo assembled scaffolds was assessed and the genomes were uploaded to the Genome OnLine Databases (GOLD) (Mukherjee, S. et al. 2021) platform. There, the assembled scaffolds were annotated (**Figure 2**) and analyzed using tools within the Integrated Microbial Genomes (IMG) and Joint Genome Institute (JGI) platforms, a manually curated genome project collection (Markowitz, V. M., et al. 2012) (https://gold.jgi.doe.gov/) (**Table 1 and Supp Tables S0, S1**).

**Figure 2:**
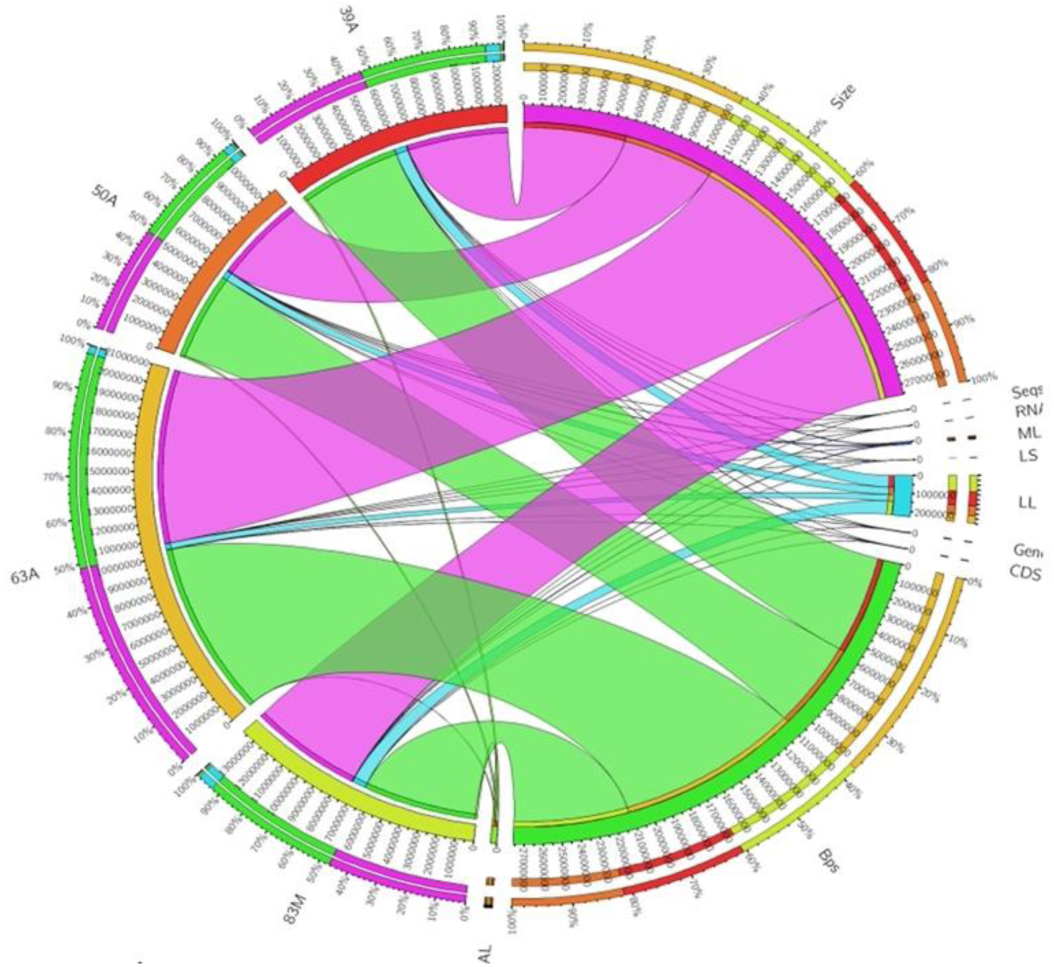
Genome Structural genome annotation Summarized the structural annotation of the four strains (63A, 50A, 39A & _19_ 83M), created using circos tool (Geers, A. U., 2021). The Genomes size (size), number of contigs (seqs), number of base pairs (Bps), Coding region (CDS), Longest length sequence (LL), Shortest length sequences (LS), Average length (AL), Median length, (ML), number of RNAs (RN), are shown as circular rings mapped back onto the strains. The inner ring presented these features in numeric, and the out rings showed them in percentages. The figure is a visual aid summary of the structural annotation of genomes features.

### BGC detection and secondary metabolite analysis

BGCs were identified using the IMG pipeline (**Figure 3**). In addition, BGCs were further identified using the antiSMASH standalone web server version 6.1.1 (Stover CK. et al. 2011). The analysis for encoded secondary metabolites or natural products was performed using the database of Natural Product Domain Seeker (NaPDoS2) version NaPDoS2_v13b (Klau, L. J., et al. 2022). In NaPDoS2, both KS and C domains were selected with default parameters of “BLASTP stringency e-value of 1e-8” and amino acid length of 200. Nucleotide and amino acid sequence files were used as inputs for these databases, respectively. The results of NaPDoS2 were used to create the phylogenetic tree of BGCs types found in our strains of genomes, and the trees are displayed using the Interactive Tree Of Life (version iTOL v5, https://itol.embl.de) (Letunic, I., & Bork, P. 2021).

**Figure 3:**
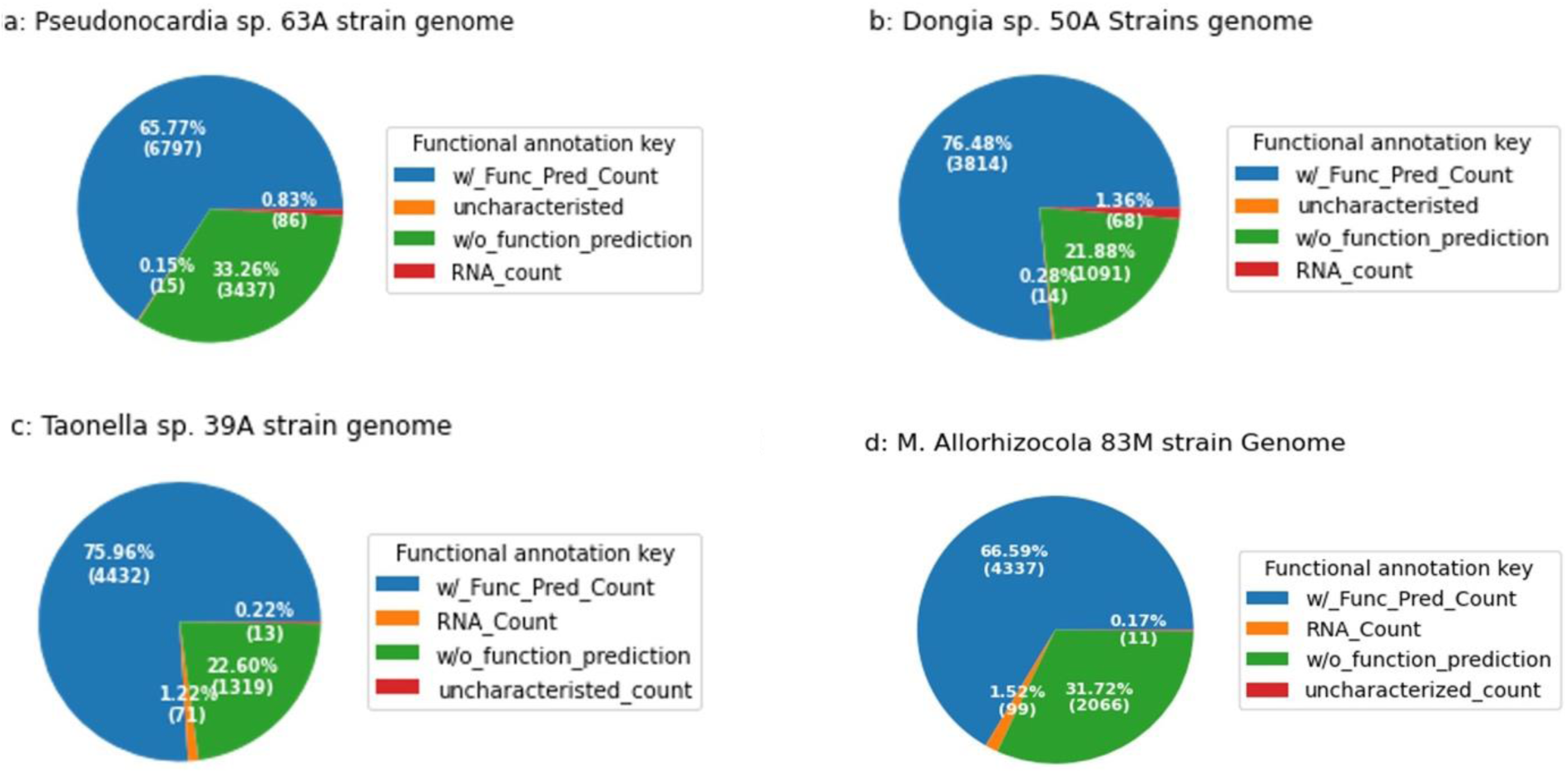
Pie charts depicting genes with predicted functions and those without predicted functions. The number of RNA’s counts and the uncharacterized number of genes in the genomes: (3а) strain 63A, (3b) 50A, (3c) 39A, and (3d) 83m. The figure shows that more significant parts of these genomes were assigned to functions (shown in blue) compared to genes with no given function (green). The other colors (red, orange) make up the RNA and uncharacterized parts of the genomes

## Results

### Genome assembly, annotation, and BCGs detection results

This study analyzes four novel bacterial strains isolated from the phyllite rocks lining the tunnel of the 17th Ledge at the SURF. After sequencing, the novel genomes were reconstructed into four drafted genomes. The quality of the de novo drafted genome assembly was assessed, and downstream analyses were performed to delimit the structural features and predicted genes/protein functions.

### Structural and functional annotations of strain 63A, 50A, 39A, and 83M

The assembled strain 63A has a genome size of 10.3 Mbp (10321584 bp), comprised of 605 scaffolds, including 581 scaffolds with and 24 without gene contents, and a total of 10335 genes in this genome. There are 10234 (99.02%) protein-coding regions (CDS) in the 581 scaffolds and 86 (0.83%) RNAs of different types, including 8 rRNAs of the subtypes made up of 3 5S, 4 16S, and 23S; 61 tRNAs, and 17 other types of RNAs such as noncoding and tmRNA). Furthermore, it contained 0.15% uncharacterized, which are mostly regulatory genes and miscellaneous features (**Supp Tables S0, S1**).

Functionally, 65.77% of CDS of this strain (63A) have predicted functions, 33.26% have no predicted functions (**Figure 3a pie chart)**, and 19.56% of protein-coding genes coded for some enzymes. Various database annotations show different percentages of sequences with assigned functions. For example, the COGs database shows 66.45%, Pfam3 shows 69.85%, and TIGRfam3 shows 16.18% assigned functions. The KEGG, KO, and MetaCyc database annotations connected 19.23%, 32.96%, and 16.65%, respectively, of protein-coding genes of this strain genome to some pathways. Moreover, 2.65% of this strain’s (63A) CDS are genes coding signal peptides, and 17.9% are genes coding transmembrane proteins, an indication of a highly active organism and a possibly competition in the dwelling environment (**Figure 4a bar chart, Supp Tables S1, S2**). Additionally, analysis of clustered genes using COG, Pfam, and TIGRfam database searches shows the presence of 1844, 2335, and 1063 clusters counted, respectively. It’s also detected 23 biosynthetic gene clusters (BGCs), comprised of 517 individual genes in this strain (63A genome) (**Figure 5a: bar chart, Table 2, Supp Table S1, S2**). These cluster genes, as reported later, were further analyzed to discover what pathways or natural products they encoded.

**Figure 4.**
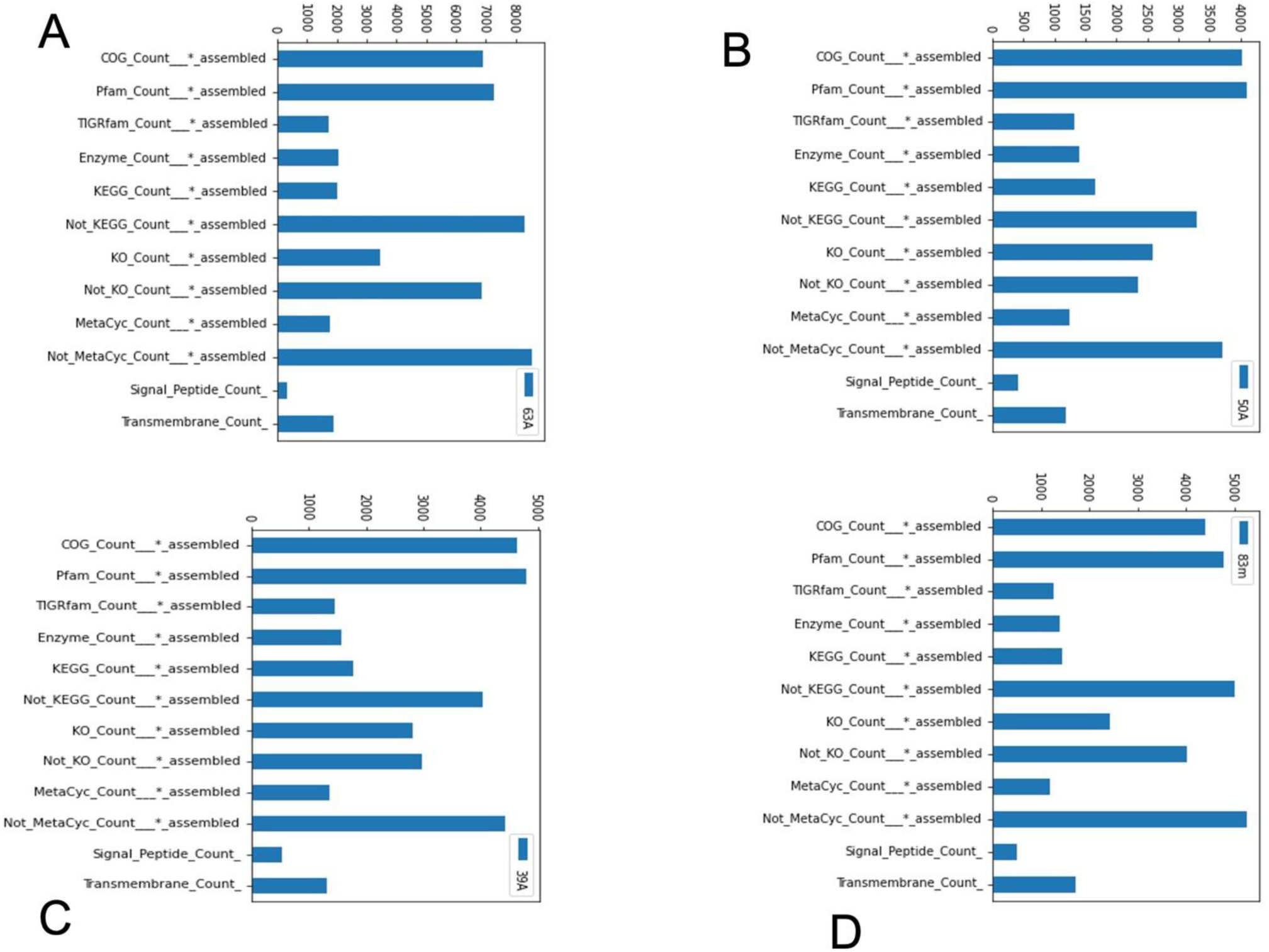
Bar plots depicting the number of proteins coding genes counts in/not in these databases from our genomes: (4a) genes count found or not found in the 63A strain genome, (4b) genes count found or not in the 50A stain, (4c) genes count found or not found in the 39A strain, (4d) genes count found or not fund in the 83m stain. Also shown are the number of proteins coding genes for signal peptides and that of transmembrane proteins.

**Figure 5:**
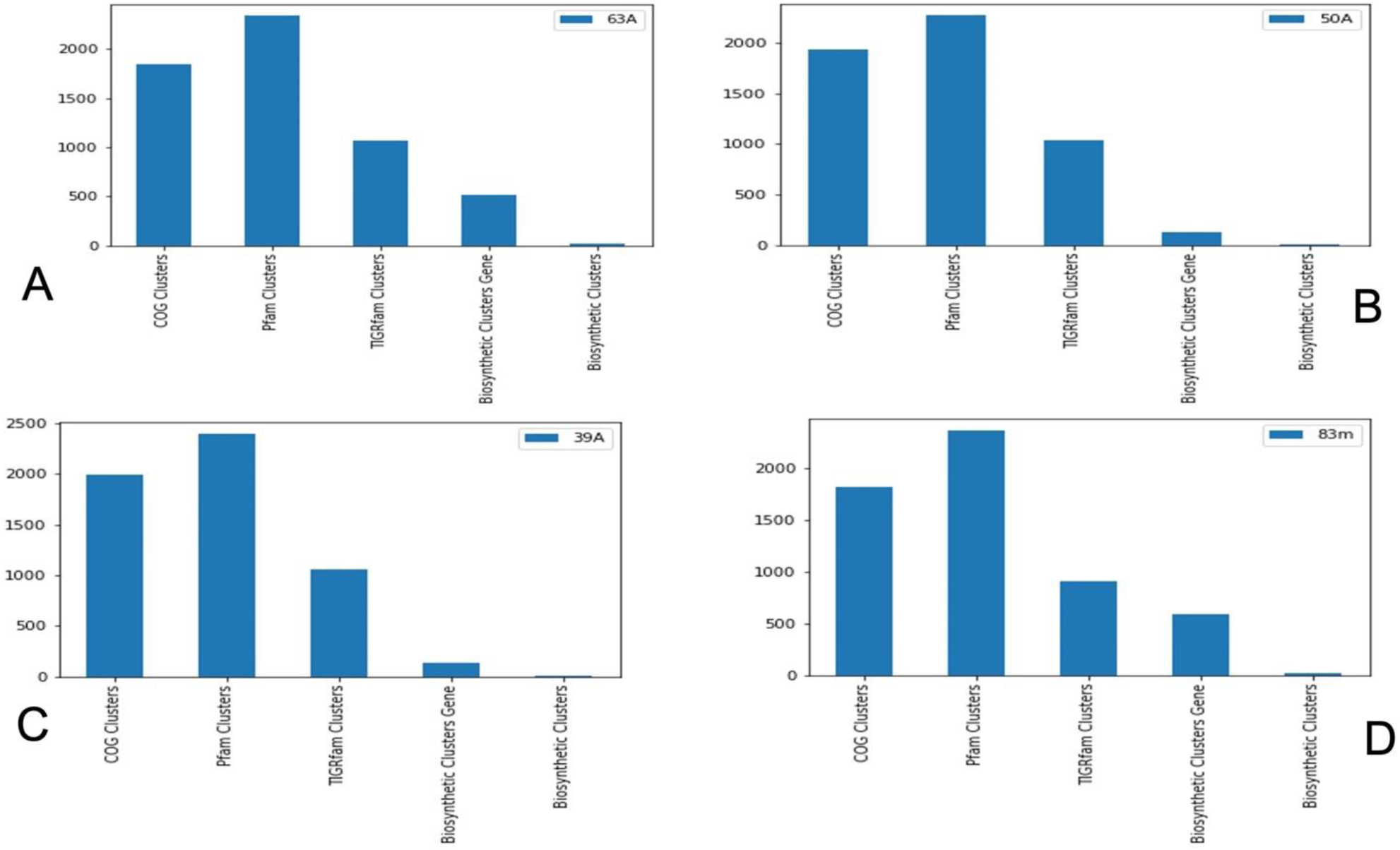
Bar plots depicting the number of clusters gene counts in COG, Pfam, and TIGRfam databases, and the total Biosynthetic Clusters counts plus the number of Biosynthetic Clusters Genes that make up each Biosynthetic Clusters in the genome of (Sa) strain 63A, (Sb) strain SOA, (Sc) strain 39A, and (5d) strain 83M, respectively. Each bar in a figure represents the number of cluster genes found in that database when the genome is annotated against the databases.

**Table 2:**
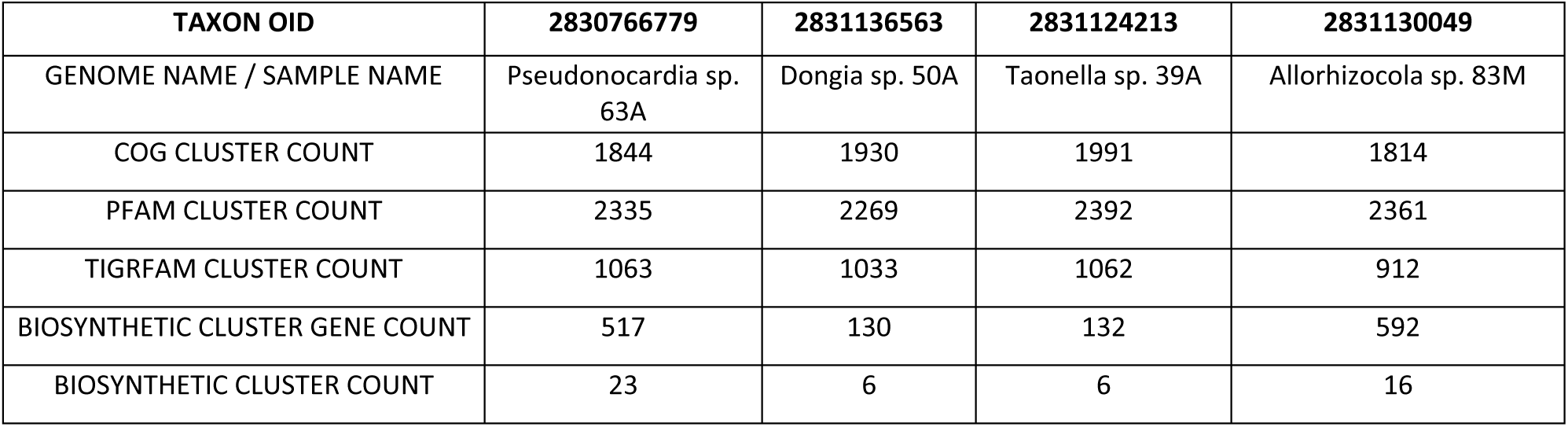
Structural annotation of genome features. These features were used to create Circle Circus Figure 2.

The assembled strain 50A resulted in 5.1 Mbp (5115127 bp) total genome size, consisting of 53 scaffolds, in which 52 of the scaffolds have genes, whereas 1 does not. There are 4987 genes in total from this 50A strain, and the genome contains 98.36% protein-coding (CDS) genes. There are 68 (1.36%) other RNA counts, including subtypes of rRNA (5S, 16S, 23S), 54 tRNAs, and 11 other RNAs types such as noncoding. The genome has 0.28% of uncharacterized genes that are regulatory and miscellaneous (**Figure 2 Circos, Table 1, Supp Tables S0, S1**).

Functional annotation assigned 76.48% of this s*train’s* CDS predicted functions and 21.88% with no predicted functions (**Figure 3b pie chart**). Additionally, 27.57% of the genome’s protein-coding genes are assigned enzyme functions. Various database annotation results show different percentages of sequences with functions assigned. For example, 80.35% of the protein-coding gene counts were assigned COG3, 81.95% Pfam3, and 25.95% TIGRfam3. The KEGG, KO, and MetaCyc databases show 32.77%, 51.59%, and 24.56% percentages of protein-coding genes connected to pathways in the respective order. Conversely, these databases also found 65.59%, 46.76%, and 73.79% of the protein-coding genes not connected to pathways. Moreover, 7.84% of protein-coding genes code signal peptides and 23.4% code transmembrane proteins in this 50A strain’s genome (**Figure 4b bar chart, Supp Tables S1, S3**). Cluster gene annotation using COG, Pfams, and TIGRfam, respectively, shows 1930, 2269, and 1033 gene cluster counts in that order. The number of biosynthetic gene clusters detected in this genome is 6, comprised of 130 individual gene counts. (**Figure 5b bar chart, Supp Tables S1, S3**). This number of BGCs is lower compared to strain 63A, although the two strains were collected from the same spot, indicating they are probably capable of different activities in the same extreme niche.

The assembled strain 39A produced a total genome size of 5.7 Mbp (5738972 bp), consisting of 49 scaffolds, in which 48 out of 49 contain genes, and 1 scaffold with no genes. The number of genes in this genome totaled 5835. There are 5751 (98.56%) protein-coding genes (CDS) in the 48 scaffolds with genes and 71 (0.22%) RNAs gene counts, including 7 rRNAs subtypes 5S, 16S rRNA, 23S rRNAs, 55 tRNAs counts, and 9 other types of RNAs. There are 0.22% uncharacterized genes in this assembled strain, which are regulatory and miscellaneous (**Figure 2 Circos, Table 1, Supp Tables S0, S1**). Functional annotation results indicate that 75.96% of the CDS have predicted functions, 22.6% lack predicted functions (**Figure 3c pie chart**), and 26.46% are protein-coding genes with enzyme-coding ability. Furthermore, 79.3% of protein-coding gene counts have assigned COG3, and that number is 81.94% for Pfam3, and 24.54% for TIGRfam3. The KEGG, KO, and MetaCyc protein-coding genes connected to pathways in these databases are 29.87%, 50.73%, and 23.05%, respectively. On the other hand, the proportion of protein-coding genes unconnected to pathways in each of these databases is 68.69% for KEGG, 50.73% for KO, and 75.51% for MetaCyc. In this 39A strain, 8.53% of protein-coding genes code for signal peptides, and 22.07% of genes code for transmembrane proteins. (**Figure 4c bar chart, Supp Tables S1, S4**). Annotation for clusters of genes show 1991 COG Clusters, 2392 Pfam Clusters, and 1062 TIGRfam Clusters. There are 6 biosynthetic gene clusters comprised of 132 individual genes in this genome (**Figure 5c bar chart, Supp Tables S1, S4**), which is also lesser than 63A strain’s BGC

Finally, the assembled 83M strain has 6.4 Mbp (6404555 bp) genome size made up of 64 scaffolds; 58 sequences have genes, and 6 scaffolds have no gene contents. The total number of genes in this assembly is 6513, with 6403 (98.31%) protein-coding genes (CDS) found in the 58 scaffolds and 99 (1.52%) RNAs genes counts included 8 rRNAs subtypes 5S, 16S, 23S, 85 tRNAs, and 6 other types of RNAs. In addition, the assembly is made up of 0.17% gene counts of uncharacterized regulatory and miscellaneous features (**Figure 2 Circos, Table 1, Supp Tables S0, S1**).

According to the functional annotation results, 66.59% of the CDS sequences of this genome have predicted functions, 31.72% have no predicted functions, and 21.02% of protein-coding genes have assigned enzyme functions (**Figure 3d pie chart**). In this strain, 67.25%, 72.96%, and 19.05% of the protein-coding genes are assigned COG3, Pfam3, and TIGRfams in the listed order. There is 21.74%, 36.77%, and 17.83% of these protein-coding genes connected to pathways and 76.57%, 61.54%, 80.49% not connected to any pathways in KEGG, KO, and MetaCyc databases, respectively. In addition, 7.5% of protein-coding genes code for signal peptides and 25.92% code for transmembrane proteins in this train (50A) (**Figure 4d bar chart, Supp Tables S1, S5**).

We examined clusters of genes using different databases and found 1814 COG clusters, 2,361 Pfams clusters genes, and 912 TIGRfams clusters genes. Additionally, this genome has 16 biosynthetic gene clusters, consisting of 592 biosynthetic genes (**Figure 5d, Supp Tables S1, S5**).

In the 63A, 50A, and 83M strains, we found Clustered Regularly Interspaced Short Palindromic Repeats (CRISPR), 1 per strain in the first two and 2 in the 83M strain. We did not detect any CRISPR sequences in the 39A strain (**Table 1, Supp Tables S0, S1**).

### Results of biosynthetic gene clusters (BGCs) mining and analysis using antiSMASH

Furthermore, using antiSMASH, we found many BGCs, with several of them putative-encoded natural products. Not to be confused, the BGCs reported in the **figure 4a-d bar plot** were obtained through the antiSMASH v5.0 database within the IMG Annotation Pipeline v.5.0.3. However, additional detection of BGCs was done using antiSMASH v6.1.1 standalone web server. Here, the nucleotides of protein-coding genes were used as inputs. Although the older and newer versions overlapped, several additional types of BGCs were detected in the genomes. The results include NRPS-like; identified as 100% icosalide A / icosalide B, NRP: Lipopeptide, a two-tailed unusual lipocyclopeptide antibiotic (Dose, B. et al., 2018), NRPS, Terpene, T3PKS, Ranthipeptide, T1PKS, Siderophore, Ladderane, Arylpolyene, and NAPAA (**Table 3**). A further analysis of Ketosynthase (KS) and Condensation (C) domains using the NPDoS2 resulted in several BGC-encoded domains. In the 63A strain, we found 4 KS and 15 C domain candidates. These are broken down according to the number of matches per class, including 6 LCL, 5 DCL, 2 epimerizations, 1 epimerization, and 1 starter (**Table 4**). In the 50A strain, there is only 1 type II FAS KS domain class, and no C domains were found, although the IMG pipeline detected the presence of BGCs in this strain’s genome. Also, in the 39A strain, there is only 1 ketosynthase (KS) domain of type II FAS class, and in the 83M, there are 11 KS domains of different classes, further divided into subclasses, and 27 C-Domains of various classes (**Table 5**)

**Table 3:**
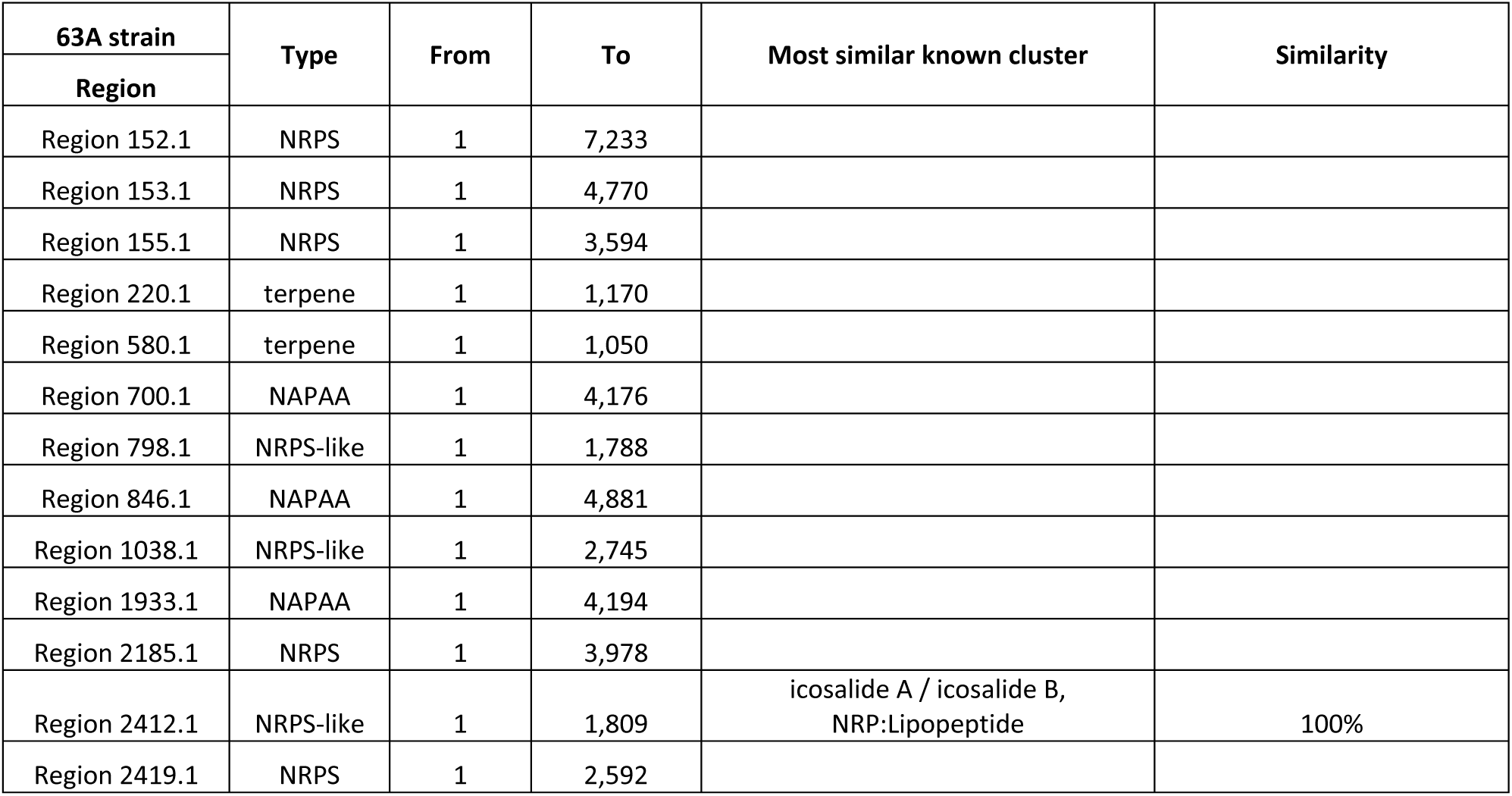
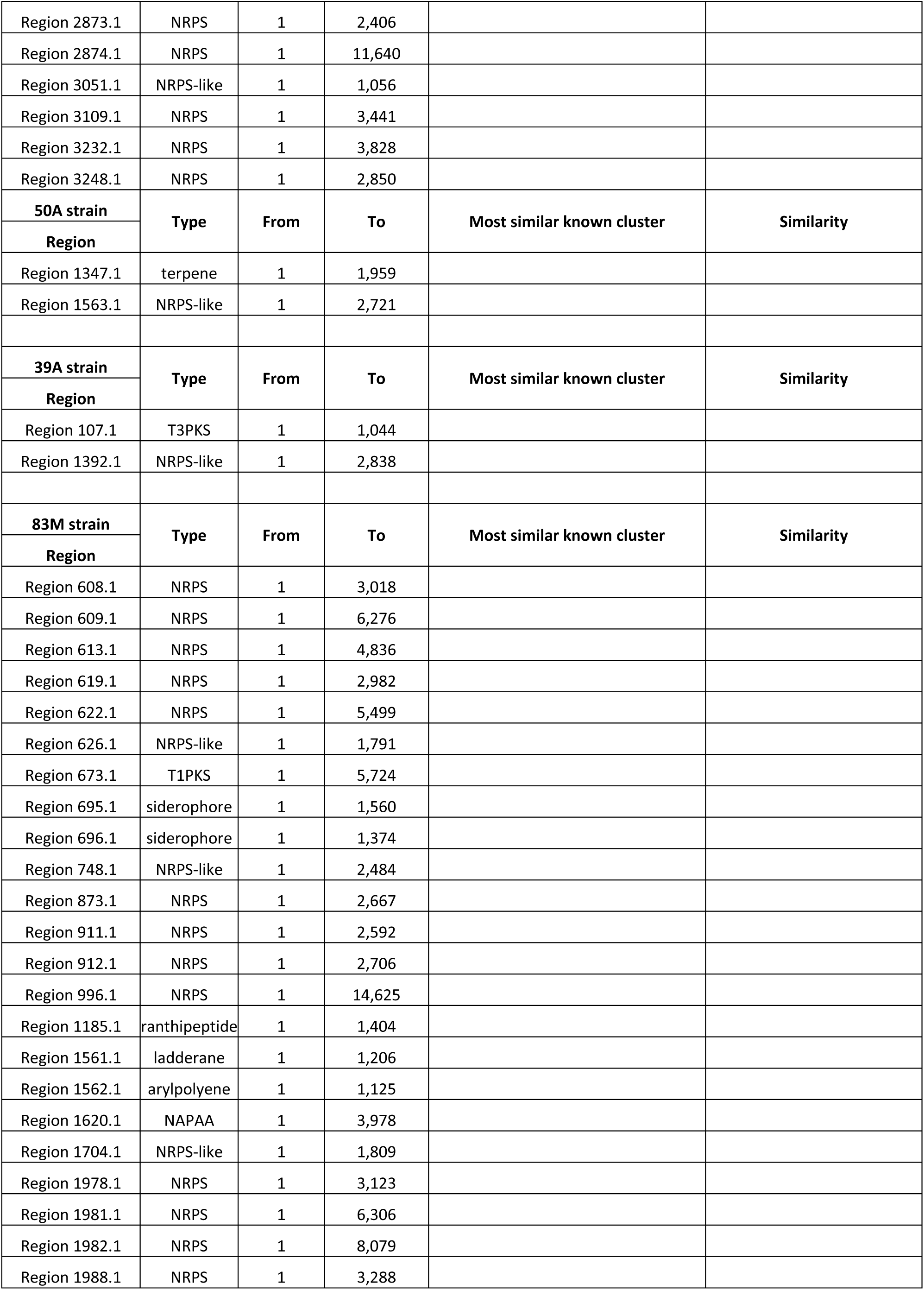
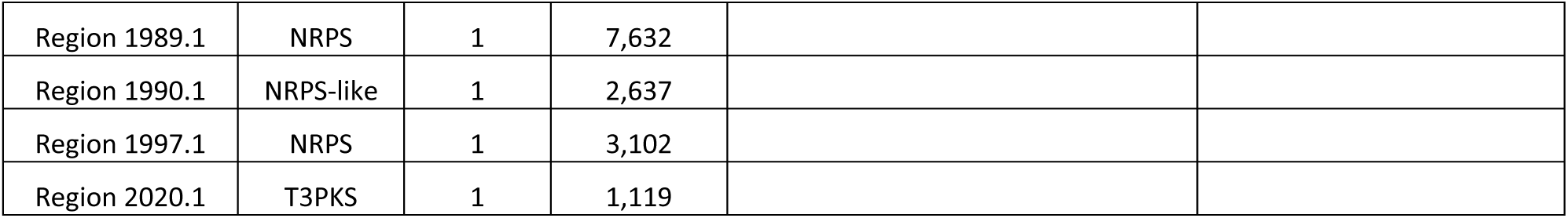
Content of several types of BGCs detected in the genomes, showing the sequence in which the type is found from start to end. The sequences are found in the supplementary files.

**Table 4:**
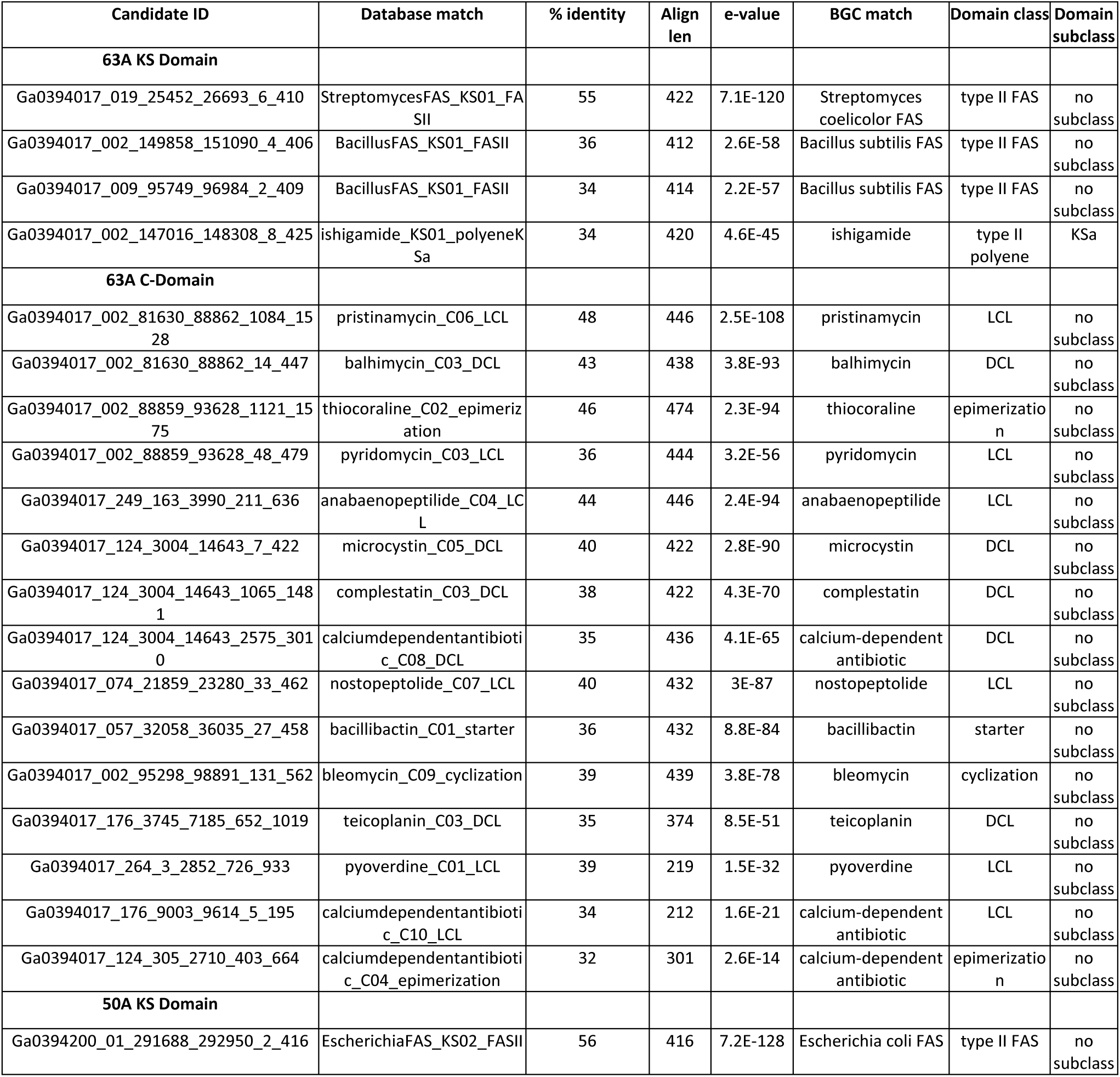
Content of the ID of candidate’s sequences in our 63A and 50A strains’ genomes that have a BGC matched, including the databases matched, percentage identity and number of amino acids matched, name of encoded BGC product, and their domains and subdomains.

**Table 5.**
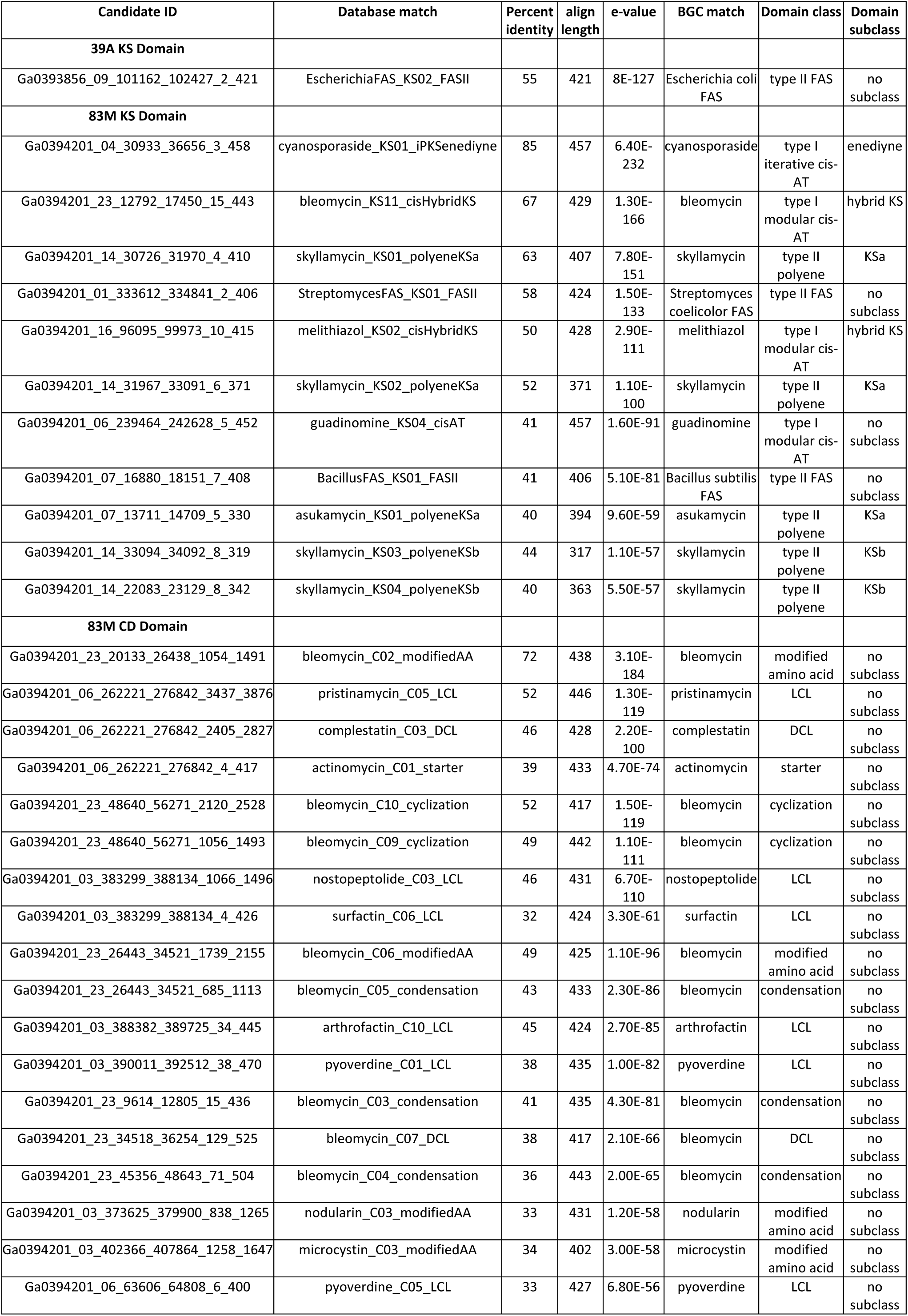
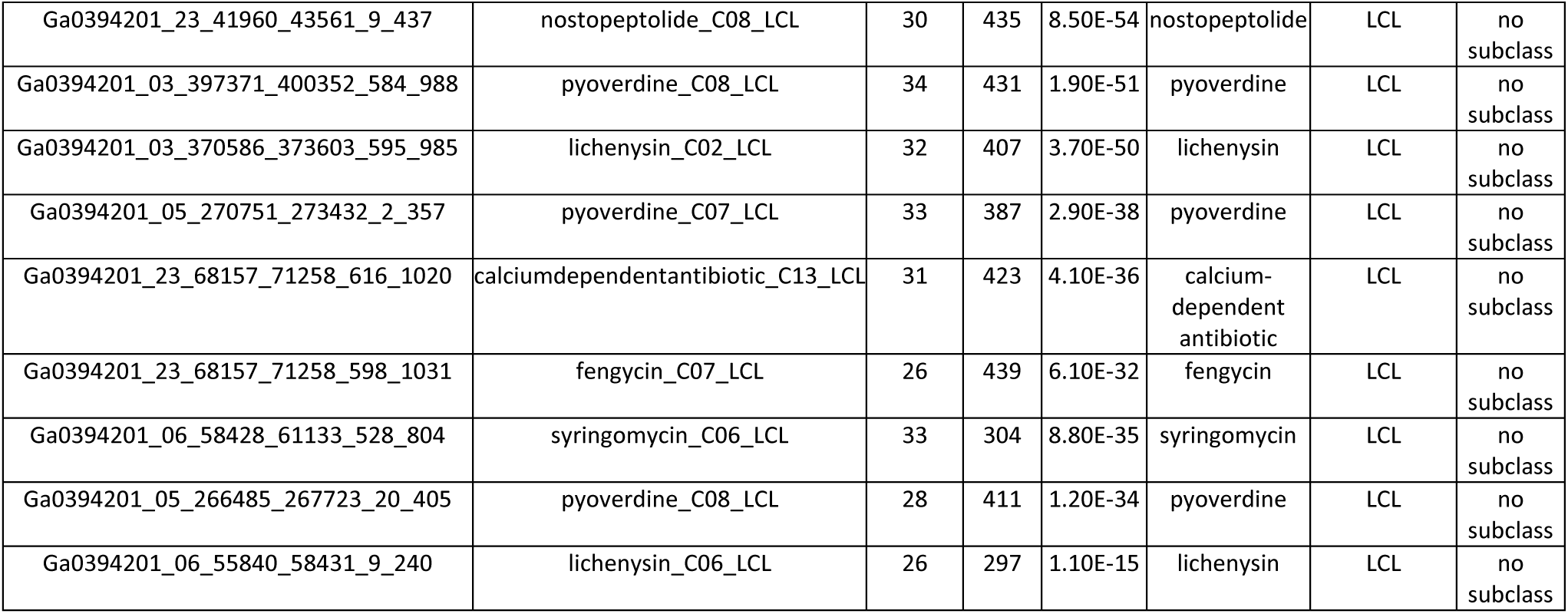
shows the content of the IDs for candidate sequences in our 39A and 83M strains’ genomes that match a BGC, including the matched databases, percentage identity, number of amino acids matched, names of encoded BGC products, and their domains and subdomains.

## Discussion

In this study, we presented the drafted de novo assembled genomes of four novel bacteria strains that were collected from the wall of a deep subsurface tunnel at the old gold mine facility. Accessing deeper subsurface resources limits the study of microbes in extreme hosting environments. In the past decade, deep subsurface sampling capability improvements and decreases in the cost of NGS have reversed this bottleneck, allowing the sampling of subsurface biospheres for bacterial biofilms. Subsurface models like the Sanford Underground Research Facility provide a conducive model to characterize the genomes of microbes dwelling in these extreme environments and allow detection of BGC genes that encode natural products important in biotechnology uses or medicine.

Although bacterial genome assembly is straightforward and made relatively easy by the lack of introns compared to eukaryotes, the contiguous arrangement makes its 80–90% protein-coding genes richer (Bohlin, J., & Pettersson, J. H. 2019). To successfully identify the features in a newly reconstructed genome requires an adequate, high-quality assembled genome. We compared our four drafted assembled genomes to the genomes of their closest reference strains, finding them to be competitive. This is indicative of a higher reconstruction quality genome. For example, the 63A strain has a genome size of 10 Mb, 10234 protein-coding genes, and 67% GC. This shows a greater similarity to its two closest genera, *Pseudonocardia acacia*, which has a genome size of 9.93 Mbp, GC of 72.3%, 66 tRNAs, and 9365 CDS (https://www.ncbi.nlm.nih.gov/genome/?term=Pseudonocardia%20acaciae[Organism]&cmd=DetailsSearch), and *Pseudonocardia spinosispora* which has a genome size of 9.5 Mbp, GC of 69.4, and 68 tRNAs (https://www.bv-brc.org/view/Genome/1123025.3).

Similarly, the 50A strain (*dongia*) has a genome size of 5.1 Mbp, and two of its closest genera, d*ongia mobilis* and d*ongia* sp., URHE0060 have a genome size between 5.0 and 6.0 Mbp, and 65 % GC (Liu, Ying., et al., 2010). The *Azospirillum* sp, the closest genera to the 39A strain, has a genome size ranging between 4.8 Mbp and 9.7 Mbp (FABIO, O. PEDROSA*, et al., 2000), putting our assembled strain in the same genome size range with *Azospirillum sp*.

Across the four genomes, the percentage of protein-coding genes with predicted functions or families is shown to be above 60% in COG and Pfam databases, but less in the TIGRfam database, the most annotated database of bacterial genomes. For example, the 63A has 66.48% COG, 69.85% Pfam, and 16.18% TIGRfam for protein-coding genes with predicted functions (**Figure 3a**). The lower hits in TIGRfam indicate that these strains are novel and haven’t been annotated thoroughly. Moreover, a smaller number of protein-coding genes is connected to pathways across the four genomes (e.g., KEGG 19.23%, MetCyc 16.65% in 63A), whereas larger percentages have no associated pathways (e.g., KEGG 79.80), MetCyc (82.37% in 63A) (**Figure 3a-d**, **Table 1, Supp Table S1**).

Additionally, the annotation indicates that each strain exhibits different expression levels of assigned functions and varying abundances of protein family domains, which explains how each strain may utilize carbon sources differently. This suggests that, despite originating from the same niche, each strain may have a distinct mechanism for carbon source utilization (**Supp Tables S2-S5**). Instead, the top abundant functions from the 63A strain are “DNA-binding transcriptional regulator”, “crR family”, “cyl-CoA dehydrogenase related to the alkylation response protein AidB”, and “MFS family permease.” These functions are for adaptive responses to changes in conditions of living and survivability strategies and are mediated by transcriptional regulator proteins (Kang, Sung-Min., et al., 2019).

The 50A, 39A, and 83M strains show a similar trend; genes needed for survival are the ones ubiquitously expressed, topping the list in all strains, although narrowly expressed at different percentage levels among the genomes. (**Supp Tables S2 – S5**)

Furthermore, active enzymatic activities were evidenced by the number of enzymes annotated per bacterium strain after removing duplicates (**Figure 6, and Supp Table S6**). The Venn diagram in **Figure 6** reveals that the strains shared a significant number of enzymes, showing common enzymes provably used for common activities by each strain and for their common environment niche. The unique enzymes found in these strains, including 134 in 63A, 81 in 50A, 65 in 39A, and 99 in 83M, may be tailored for the specific tasks of each individual bacterial strain. One of the shared 454 enzymes is the “mesaconyl-CoA hydratase β-methylmalyl-CoA dehydratase,” a ubiquitous family with functions in the central carbon metabolism pathway in bacteria (Zarzycki, J., 2008). It is possible that a further analysis of these enzymes might reveal putative activities of each strain in a biofilm environment, as we had also detected genes involved in quorum sensing pathways.

**Figure 6:**
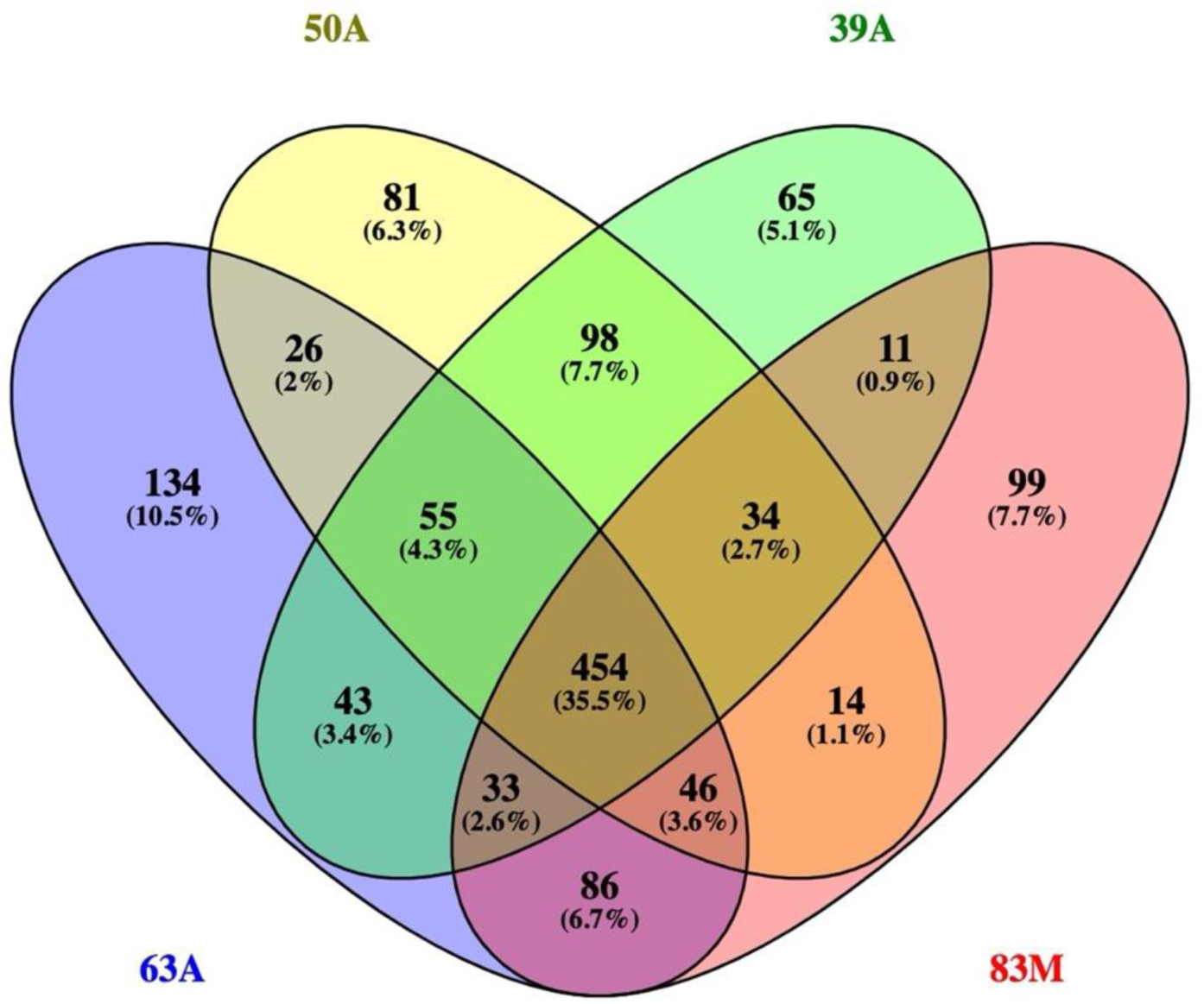
A Venn diagram of distinct EC expressed in each strain and the overlapping EC between the isolate’s strains’ genomes. 454 enzymes are present in all the strains, and 134 EC are unique to 63A, 81 EC for 50A, 65 for 39A, and 99 for 83M strain.

Furthermore, the level of horizontal gene transfer in these strains’ genomes demonstrates their genome plasticity. In strains 63A, 50A, and 39A, we found 20.75%, 24.18%, and 31.26% composition of horizontal gene transfer, respectively. However, strain 83M has a much higher horizontal transfer above 32% (**Supp table 1**). These percentages are way beyond the expected or known horizontal gene transfer or plasticity. The current documented horizontal gene transfer in bacteria can reach up to 15% (Garcia-Vallvé, S. et al., 2000), which places the first three strains at the edge of the known gene transfer spectrum, while strain 83M exhibits remarkably high horizontal gene transfer. Conversely, given the extreme environment where the samples were collected (deep subsurface old mine biofilm samples), it is possible to speculate that these transfers might be genuine. The extreme underground dwelling niche could potentially explain the higher gene transfer, as the competition for nutrients could have triggered the incorporation of horizontal genes for survival. Moreover, the high gene transfers could have been facilitated by the rapid process of biofilm formation orchestrated by extreme biospheres.

As we searched for biosynthesis gene clusters (BGCs) and important secondary metabolite-producing pathways in our genomes, we found several secondary metabolites, or natural products, important for biotechnology and medicine. Although it is still early to make a claim, the discovery of biosynthetic gene clusters (BGCs) in the genomes of our strains suggests that they have the potential to produce secondary metabolites. Further studies could potentially utilize these strains for their significant potential, and if investigated further, these strains could serve as machinery for producing natural products. For example, non-ribosomal peptides, a class of secondary metabolites with many properties, such as toxins, siderophores, pigments, or antibiotics (Martínez-Núñez et al., 2016), are found in our assembled genomes, making these bacteria strains highly important to explore more. Currently, the significance of BCGs (biosynthetic cluster genes) is expanding among the biotechnology and pharma industry, making their discovery a hot topic (Russell, A. H., & Truman A. W., 2020, Medema M. et al., 2014).

As we did the analysis using the antiSMASH (Stover CK. et al. 2011) database, we confirmed the presence of BGCs and analyzed their encoded products using the NaPDoS2 (Klau, L. J., et al. 2022) database (**Table 1, Supp S0, S1, Figures 4a-d**). Reported in **Figure 4a-d**, and **Tables 1& 3**, are BGCs that were identified in each genome through the IMG pipeline and confirmed again using a standalone antiSMASH. **Figure 7 Venn diagram** shows the number of intersected BGCs between the four genomes after removing duplicates. Excluding the “hypothetical proteins,” the three common BGCs identified are those encoding for “SAM-dependent methyltransferase,” “DNA-binding transcriptional LysR family regulator,” and “signal transduction histidine kinase.” The SAM-dependent methyltransferase is a ubiquitous cofactor enzyme that is used in many reactions; however, it played a vital role in the biosynthesis of natural products with prolific “pharmacological moieties” (Sun, Q. et al., 2021). The LysR family regulator is another prevailing enzyme in prokaryotes and a positive regulator of transcription initiation (Schell, M. A., 1993). The signal transduction histidine kinase is a multifunctional enzyme that cascades signals across cell membranes in bacteria (Mascher, T. et al., 2006). The active presence of this enzyme in all four strains might be explained by the critical functional role they play in the lives of those bacteria in that extreme environment niche.

**Figure 7:**
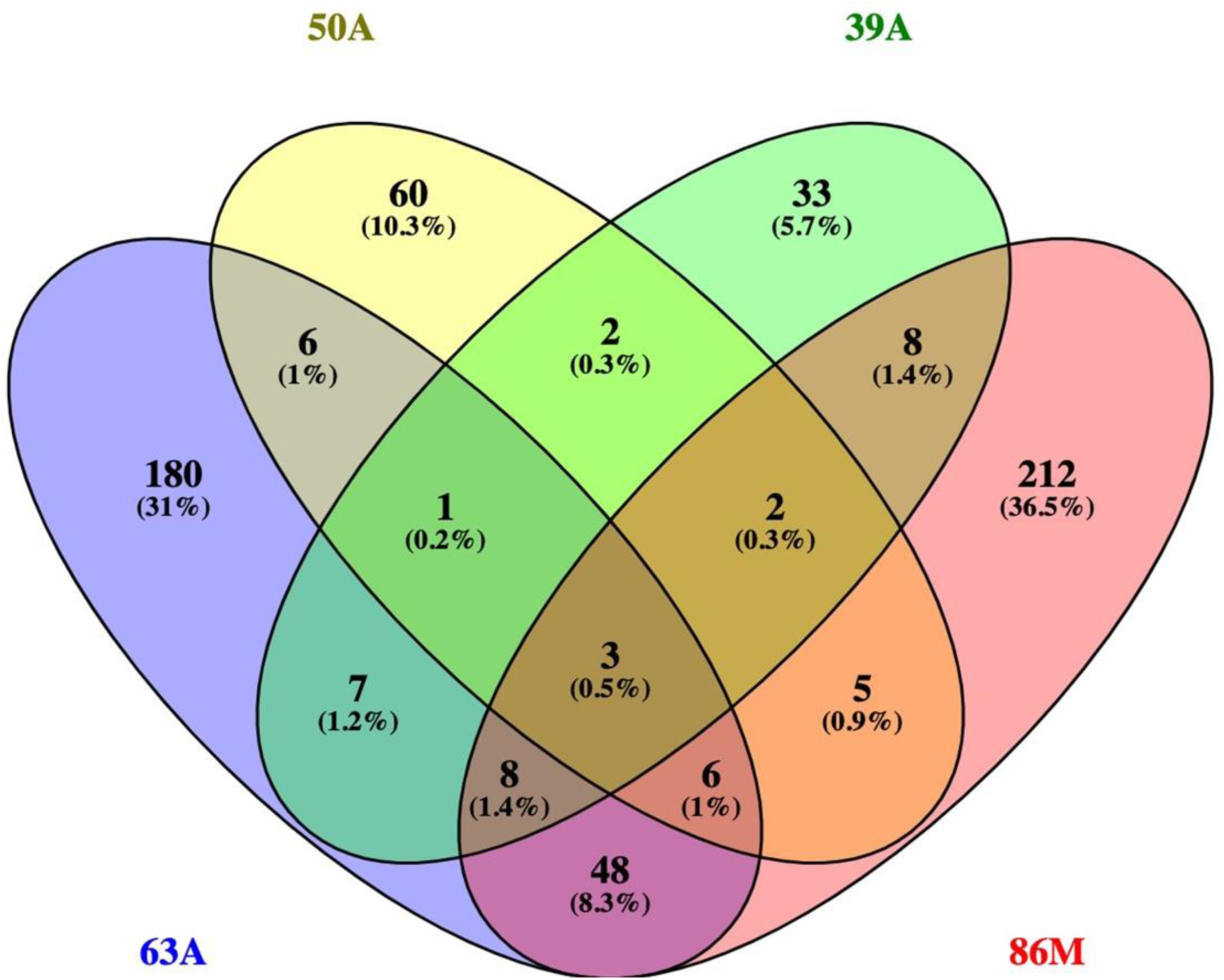
Venn diagram depicting the number of shared and unique BGCs genes detected in these novel strains’ genomes after duplicates are removed. The three common elements in “63A,” 50A, “39A,” and 83M are “SAM-dependent methyltransferase,” “DNA-binding transcriptional LysR family regulator,” and “signal transduction histidine kinase”

We analyzed the same types of BGCs using antiSMASH v6.1, a standalone web server, which confirmed the accuracy of the first pipeline finding. Details of these findings include NRPS-like 100% annotated as icosalide A / icosalide B, NRP:Lipopeptide, a two-tailed unusual lipocyclopeptide antibiotic (Dose, B. et al., 2018), terpene, NRPS, and NAPAA (**Table 3**). Lipopeptides are used as antimicrobial, antifungal, and antitumor agents (Kourmentza K. et al., 2021). We also detected NRPS-like catalyzed peptide metabolites, which are natural products for making important drugs, including vancomycin, daptomycin, and cyclosporine A (Shi J. et al., 2021). Similarly, we identified BGCs encoding antimicrobials, such as surfactin, iturin, and fengycin (Raaijmakers, J. M. et al., 2010; Raaijmakers, J. M. et al., 2010). Other natural products found include terpene and NRPS-like detected in strain 50A and annotated as AMP-binding and NTPD-like fragments. In addition to these found in the other genome strains, 83M has four types of BGCs that were unique to its genome (**Table 3**).

Assessment of mechanisms and potential functions using domain analysis shows KS and C-domains processes, including 4 Ketides Synthases and 15 Condensation domains candidates in the 63A strain. **Table 4** contains the breakdown of those domains according to the number of matches per class. One of the C domains we detected is that of epimerization, a process used to make natural products such as the drug tetracycline. The presence of cluster genes encoding these pathways in this strain suggested its mechanism for producing secondary metabolites. These findings call for further investigation to confirm and understand these systems in this strain, as well as their potential. We also found type II FAS (**Table 3**) in the 50A strain, by analyzing the KS domains; however, no C-domains class of metabolites was matched despite the previous report of BGCs in this genome from the IMG pipeline (**Figure 3b**).

In support of novelty, it is documented that lower levels of BGCs (< 85%) in the genome indicated possible novelty of the genome (Medam et al., 2015, Chen et al., 2020); thus, low percentage hits of BGCs, particularly in the 50A strain, show a possible presence of yet-to-be-discovered BGCs, although this statement is speculative and requires more evidence. In the 39A strain, there is one KS domain belonging to type II FAS (**Table 3**). Type II FAS are less understood biochemical metabolites; however, a recent study has suggested that they play a critical role in the enzymatic pathways with the help of acyl carrier protein (ACP) (Chen A. et al., 2018). Interestingly, ACP was among the top overexpressed functions in the 39A strain genome.

Besides BGCs detected in the other strains, Ranthipeptide, T1PKS, Siderophore, Ladderane, and Arylpolyene were detected only in strain 83M (**Tables 3 & 4**). Siderophore pathways are responsible for nonlinear biosynthesis and are considered megasynthases. They are related to fatty acid synthases that biosynthesize diverse natural-function small molecule polyketides (Martínez-Núñez and López S., 2016). Ranthipeptide is an emerging novel class of natural products, classified as the ribosomally synthesized, post-translationally modified peptide (RiPP) superfamily (Chen, Y. et al., 2021), and it is suggested to be involved in quorum sensing (Chen, Y. et al., 2020). Ladderane, however, is used by bacteria in anaerobic ammonium oxidation, which includes proton and hydroxide gradients control in bacteria “ammonoxosome” compartments. (Pályi, G., 2020). Arylpolyene is a natural product class that provides red pigmentation. Bacteria produce arylpolyene in abundant amounts, which functions in the uptake of UV light (Schöner, T. A. et al., 2016), thus used as a food additive and antioxidative, and has health benefits (Kong K. W. et al., 2010).

Lastly, these genomes contain Clustered Regularly Interspaced Short Palindromic Repeats (CRISPR) sequences that demonstrate the strains’ capability. Bacteria use CRISPR to switch on and off specific genes as needed in response to internal or external signaling (Redman, M., 2016). Single CRISPR was found in the 63A and 50A strains, and two in the 83M, whereas no CRISPR was found in the 39A strain (**Table 1, Supp Tables S0, S1**). The three strains with CRISPR are likely utilizing these systems to respond to internal or external signals related to the demands of their hosting environment; however, this warrants further investigation.

### Conclusion

Generally, sampling deep subsurface environments like SURF could enhance our understanding of the microbiome found in underground areas. Deep hosting biospheres provide the right conditions that lead to microbes’ capabilities not found in other environments, such as the utility of different forms or types of carbon sources, biofilm formation, and development of capabilities to produce important secondary metabolites. These conditions make the extreme hosting environment a fertile place for the possible discovery of novel genes or pathways useful for new biotechnology development.

In this study we use the Sanford Underground Research Facility (SURF), an old gold mine turned into a research laboratory, as a model of a subsurface environment to obtain and characterize four novel strains of bacteria by sequencing, reconstructing, and analyzing their genomes. This resulted in four novel genomes of varied sizes or lengths, annotated against various databases using the IMG annotation pipelines, antiSMASH, and the NaPDoS web servers, respectively. Genome annotation shows different structural and functional genome properties reported in the results sections. BCGS mining shows several types of secondary metabolites, or compounds encoded by the BGCs clusters, such as polyketides (PKS, PKS III, Ketosynthases domain-KS), non-ribosomal peptides (condensation domain, NRPS), and terpenoids (terpenes I, polyenes BCG type II) in the genomes. These secondary metabolites, or natural products, are used in many ways, including medicines and biotechnology products. Therefore, based on our findings, these novel genomes may serve as valuable resources for the functions discussed above. Furthermore, our finding highlighted the importance of subsurface environments in enhancing bacteria capabilities, turning them into natural product producers.

In all, extreme hosting niches are a beneficial model for studying and finding novel strains of bacteria useful in biotechnology. We believe the four drafted novel genomes could add value to a future understanding of a deep subsurface microbiome and the discovery of enhanced strains of bacteria. Moreover, this research will enhance our understanding of how microbes from endogenous or surface sources colonize deep subsurface habitats, develop various pathway capabilities, such as encoding secondary metabolites of medical importance, and how these systems may lead to the discovery of new biotechnology applications and developments.

## Data availability

Data Records Under the BioProject ID: PRJNA551088, raw data sequences from the four isolated and sequenced microbial (Allorhizocola sp 83M, Taonella-related sp. 39A, Dongia-related sp. 50A, and Pseudonocardia sp. 63A), were deposited to the National Center for Biotechnology Information (NCBI) Sequence Read Archive (SRA) as SRR9595710, SRR9600320, SRR9600318, SRR9600319; Accession number: SAMN12138386, SAMN12139076, SAMN12139075, and SAMN12139074, respectively. The four raw NGS datasets described in this manuscript can be downloaded from the NCBI’s SRA database using the accession numbers provided in **Table 2**. On the other hand, detailed instructions for downloading datasets from NCBI SRA are supplied on the NCBI website: https://www.ncbi.nlm.nih.gov/sra/ for further help. Additionally, downstream analysis of this data is also provided on the GOLD/IMG/EMR website. This included the four contigs/scaffolds, and their annotation results obtained from the JGI (Joint Genome Institute) pipeline annotation for contigs/scaffolds; see **Table 4**.

## Funding Statement

Research reported in this publication was supported by the South Dakota Biomedical Research Infrastructure Network (SD BRIN (Biomedical Research Infrastructure Network)) through an Institutional Development Award (IDeA) from the National Institute of General Medical Sciences of the National Institutes of Health under grant number P20GM103443.

Also, supported by a fellowship to (DA) by the USD Neuroscience, Nanotechnology, and Networks (USD-N3) program through a grant from NSF (NATIONAL SCIENCE FOUNDATION) (DGE-1633213), and supported by the National Science Foundation/Experimental Program to Stimulate Competitive Research (EPSCoR) Grant IIA 1355423. The BioSNTR grant UP1700139A partially funded this work from the National Science Foundation/Experimental Program to Stimulate Competitive Research (EPSCoR) Grant OIA-1849206 awarded to Gilbert Ustad and the Institutional Development Award (IDeA) from the National Institute of General Medical Sciences of the National Institutes of Health P20GM103443 (V.C.Huber). The content expressed here is solely the responsibility of the authors. It does not necessarily represent the official views of the National Institutes of Health, NSF, USD-Neuroscience, Nanotechnology and Networks program, or SCORE Track I.

## Acknowledgements

West Core performed the collection of samples and the NGS sequencing at Black Hills State University (BHSU). The Bioinformatic Core at the University of South Dakota (USD) supported the data analysis.

## Author Contributions

**DA** did the data analysis and wrote the first and final draft, as well as reviewed the manuscript.

**JZ** helped review and edit the final manuscript.

**SA** built the figure in BioRender and helped review and edit the final manuscript.

**OG** sequenced the genomes and drafted the sequencing method section.

**DB** collected the samples, isolated and identified the bacteria strains using 16S, wrote some of the manuscript introduction, and helped review the final manuscript.

**VG** helped review and edit the final manuscript.

**EG** directed the experimental design and data analysis and helped in the review of the final manuscript.

## Conflict of Interest

The authors have declared no conflict of interest.

